# Left prefrontal impact links subthalamic stimulation with depressive symptoms

**DOI:** 10.1101/665976

**Authors:** Friederike Irmen, Andreas Horn, Philip Mosley, Alistair Perry, Jan Niklas Petry-Schmelzer, Haidar S. Dafsari, Michael Barbe, Veerle Visser-Vandewalle, Gerd-Helge Schneider, Ningfei Li, Dorothee Kübler, Gregor Wenzel, Andrea Kühn

## Abstract

**Objective:** Subthalamic nucleus deep brain stimulation (STN-DBS) in Parkinson’s Disease (PD) not only stimulates focal target structures but also affects distributed brain networks. The impact this network modulation has on *non-motor* DBS effects is not well characterized. By focusing on the affective domain, we systematically investigate the impact of electrode placement and associated structural connectivity on changes in depressive symptoms following STN-DBS which have been reported to improve, worsen or remain unchanged.

**Methods:** Depressive symptoms before and after STN-DBS surgery were documented in 116 PD patients from three DBS centers (Berlin, Queensland, Cologne). Based on individual electrode reconstructions, the volumes of tissue activated (VTA) were estimated and combined with normative connectome data to identify structural connections passing through VTAs. Berlin and Queensland cohorts formed a training and cross-validation dataset used to identify structural connectivity explaining change in depressive symptoms. The Cologne data served as test-set for which depressive symptom change was predicted.

**Results:** Structural connectivity was linked to depressive symptom change under STN-DBS. An optimal connectivity map trained on the Berlin cohort could predict changes in depressive symptoms in Queensland patients and vice versa. Furthermore, the joint training-set map predicted changes in depressive symptoms in the independent test-set. Worsening of depressive symptoms was associated with left prefrontal connectivity.

**Interpretation:** Fibers linking the STN electrode with left prefrontal areas predicted worsening of depressive symptoms. Our results suggest that for the left STN-DBS lead, placement impacting fibers to left prefrontal areas should be avoided to maximize improvement of depressive symptoms.

## Introduction

Deep brain stimulation (DBS) of the subthalamic nucleus (STN) provides relief of motor symptoms in movement disorders such as Parkinson’s disease (PD) by exerting influence on focal target structures and distributed brain networks^1^. A strong relationship between connectivity profiles of DBS electrodes and clinical improvement has been shown in PD^2^ and recently in patients with obsessive compulsive disorder^3^. While Horn *et al.*, 2017 describe the structural connectivity profile associated with *motor* improvement in PD, little is known on how structural connectivity impacts *non-motor* DBS effects. Currently accepted theoretical frameworks postulate that non-motor symptoms following DBS stimulation of basal ganglia targets originate from the modulation of overlapping cortex-basal ganglia motor and non-motor loops^4^. In PD, non-motor DBS-effects have been described in various domains including autonomic function, sleep, cognition and mood^5–9^. In the affective domain, in addition to postoperative hypomania^10^, acute depression can also be a side effect of STN-DBS in PD patients^11,12^ with a prevalence of about 20-25%^13^ despite slight improvement after 6 months. Interestingly, STN-DBS has been reported to *improve, worsen* or to have *no effect*^9^ on symptoms of depression or anxiety. However, unlike mania, postoperative depressive symptoms have rarely been associated with sensorimotor STN stimulation itself but rather with too fast tapering of dopaminergic medication^14^ and stimulation of more ventral STN territory or even zona incerta stimulation^13,15,16^. Indeed, the precise local placement of DBS electrodes impacts non-motor DBS effects^17,18^ and modulation of remote brain regions involved in affective processing might play a crucial role on how affective symptoms develop after surgery. In this study, we follow up on the research of Horn *et al.*, 2017 by investigating the impact of electrode placement and associated structural connectivity on changes in depressive symptoms following STN-DBS. To this end, we reconstructed electrode placement in 80 PD patients from two international DBS centers and estimated their structural connectivity profiles using age-matched normative connectome that was acquired in a different sample of PD-patients. Based on these connectivity profiles, we calculated models that could explain and cross-predict worsening or improvement in depressive symptoms as measured with the Beck Depression Inventory-2^nd^ Edition (BDI-II)^19^. Finally, we validated these models using a testing dataset of 36 PD patients from a third DBS center.

## Materials and methods

### Patient cohorts and imaging

A total of 121 patients from three DBS centers (Berlin [BER]: n=32; Queensland [QU]: n=49; Cologne [CGN]: n=40) were included in this retrospective study (age 62±9 years, 43 women; detailed clinical data in Supplement 1). Data from BER and QU were used to form the *training* and *cross-validation datasets* to identify structural connectivity predicting mood changes after DBS surgery. Data from the CGN was used as a *test dataset* to validate the established model. Five patients were excluded from the analyses for the following reasons: One patient (QU) due to incomplete data, two patients (CGN) due to unilateral VIM (instead of STN) stimulation, and two patients (CGN) due to clinically relevant psychiatric comorbidities that were pharmacologically treated in the same timeframe in which BDI-II changes were assessed. Table 1 summarizes the sample characteristics of the final cohort (n=116).

**Table 1:** Sample characteristics.

|  | BERLIN |  | QUEENSLAND |  | COLOGNE |  | TOTAL |  |
| --- | --- | --- | --- | --- | --- | --- | --- | --- |
| **N** | 32 |  | 48 |  | 36 |  | 116 |  |
| **SEX** | 10f | 22m | 15f | 33m | 18f | 18m | 43f | 73m |
|  | *M* | *SD* | *M* | *SD* | *M* | *SD* | *M* | *SD* |
| **AGE (YRS)** | 61 | 9 | 62 | 9 | 62 | 8 | 62 | 9 |
| **DISEASE DURATION (YRS)** | 10 | 5 | 8 | 4 | 10 | 3 | 9 | 4 |
| **MONTHS POST SURGERY** | 12 |  | 6 |  | 6 |  | 7 |  |
| **BDI-II (BASELINE)** | 11.56 | 6.15 | 11.06 | 4.69 | 7.00 | 4.22 | 9.94 | 5.41 |
| **BDI-II (POSTOP)** | 11.56 | 7.33 | 8.45* | 5.60 | 7.00 | 5.76 | 8.96 | 6.43 |
| **UPDRS-III (BASELINE, MED ON)** | 20.78 | 10.16 | 37.46 | 15.26 | 17.78 | 9.75 | 26.75 | 15.45 |
| **UPDRS-III (ON DBS, MED ON)** | 19.26 | 13.55 | 33.95 | 12.94 | 17.00 | 8.79 | 24.85 | 14.44 |
| **LEDD-REDUCTION (%)** | 46.06* | 40.06 | 68.98* | 22.75 | 48.27* | 18.66 | 56.32* | 29.70 |

BDI-II – Delta change in Beck’s depression inventory (Baseline = pre; Postop = post DBS surgery); UPDRS-III –Unified Parkinson’s disease rating scale III (Baseline = pre; Postop = post DBS surgery ON Medication); LEDD – Levodopa-equivalent daily dosage; M – mean; SD – Standard deviation; \*significant change compared to baseline

All patients underwent stereotactic DBS surgery for treatment of PD (surgical methods described elsewhere^20^) and received bilateral DBS electrodes (n=42 model 3389 Medtronic, Minneapolis, MN; n=31 Boston Scientific Vercise; n=36 Boston Scientific Vercise Cartesia Directional; n=7 St Jude Infinity Directional model 6172). Structural abnormalities were excluded using preoperative MRI. Clinically significant psychiatric symptomatology and cognitive deficits (defined as deficient performance in Mini-Mental State Examination score or multidomain deficits in neuropsychological tests such as features of PD dementia) were assessed by psychiatric evaluation and neuropsychological testing as exclusion criteria prior to surgery. In all patients, lead placement was validated using microelectrode recordings during surgery, intraoperative macrostimulation and postoperative imaging. In all three DBS centers, quality of pre- and postoperative MRI and CT data was controlled by meticulous visual inspection during stereotactic planning by an interdisciplinary team of neuroradiologists, neurologists and neurosurgeons. In case of even slight movement artefacts that reduced image quality in a patient, image acquisition was repeated under general anaesthesia before surgery. Depressive symptoms were recorded prior to surgery and postoperatively (after 7.56±2.9 months [i.e. M±SD throughout the paper], when DBS settings had been titrated intensively and stable settings have been reached) using BDI-II (cut-off values 0-13:minimal depression; 14-19:mild depression; 20-28:moderate depression; >29:severe depression). The BDI-II has been validated as a reliable tool for assessment of depressive symptoms in PD despite its disadvantage of potentially biased responses due to face validity. However, it has the practical advantage of an easy and fast administration and is therefore routinely acquired in DBS care units, and widely used in studies assessing DBS effects^21^. To control for covariates, sex, levodopa equivalent daily dosage (LEDD), LEDD of dopamine agonists (LEDD-DA), and Unified Parkinson’s Disease Rating Scale Part III (UPDRS-III) ON medication were recorded preoperatively and postoperatively ON DBS in all patients. UPDRS-III data OFF medication was only available for the BER cohort. The study was approved by the local ethics committee at each site and carried out in accordance with the Declaration of Helsinki.

### Localization of DBS electrodes

DBS electrodes were localized using the advanced processing pipeline^20^ in Lead-DBS (www.lead-dbs.org^20^). In short, postoperative CT or MRI were linearly coregistered to preoperative MRI using advanced normalization tools (ANTs; stnava.github.io/ANTs/). Coregistrations were inspected and refined if needed. A brainshift correction step was applied as implemented in Lead-DBS. All preoperative volumes were used to estimate a precise multispectral normalization to ICBM 2009b NLIN asymmetric (“MNI”) space applying the ANTs SyN Diffeomorphic Mapping using the preset “effective: low variance default + subcortical refinement”. In some patients where this strategy failed, a multispectral implementation of the Unified Segmentation approach implemented in Statistical Parametric Mapping software (SPM12;http://www.fil.ion.ucl.ac.uk/spm) was applied. These two methods are available as pre-sets in Lead-DBS and were top-performers to segment the STN with precision comparable to manual expert segmentations in a recent comparative study^22^. DBS contacts were automatically pre-reconstructed using PaCER or the TRAC/CORE approach and manually refined if needed^20^. For segmented leads, the orientation of electrode segments was reconstructed using the Directional Orientation Detection (DiODe) algorithm^23^.

### Volume of Tissue Activated and connectivity estimation

Fig. 1 provides an overview of the methodology. The volume of tissue activated (VTA) was calculated using default settings in Lead-DBS applying a Finite Element Method (FEM)-based model^2^. This model estimates the E-field (i.e. the gradient distribution of the electrical charge in space measured in V/mm) on a tetrahedral mesh that differentiates four compartments (grey and white matter, electrode contacts and insulation). Grey matter was defined by key structures (STN, internal and external pallidum, red nucleus) of the DISTAL atlas^24^. The resulting gradient vector magnitude was thresholded at a heuristic value of 0.2 V/mm to generate the VTA^2^.

**Figure 1:**
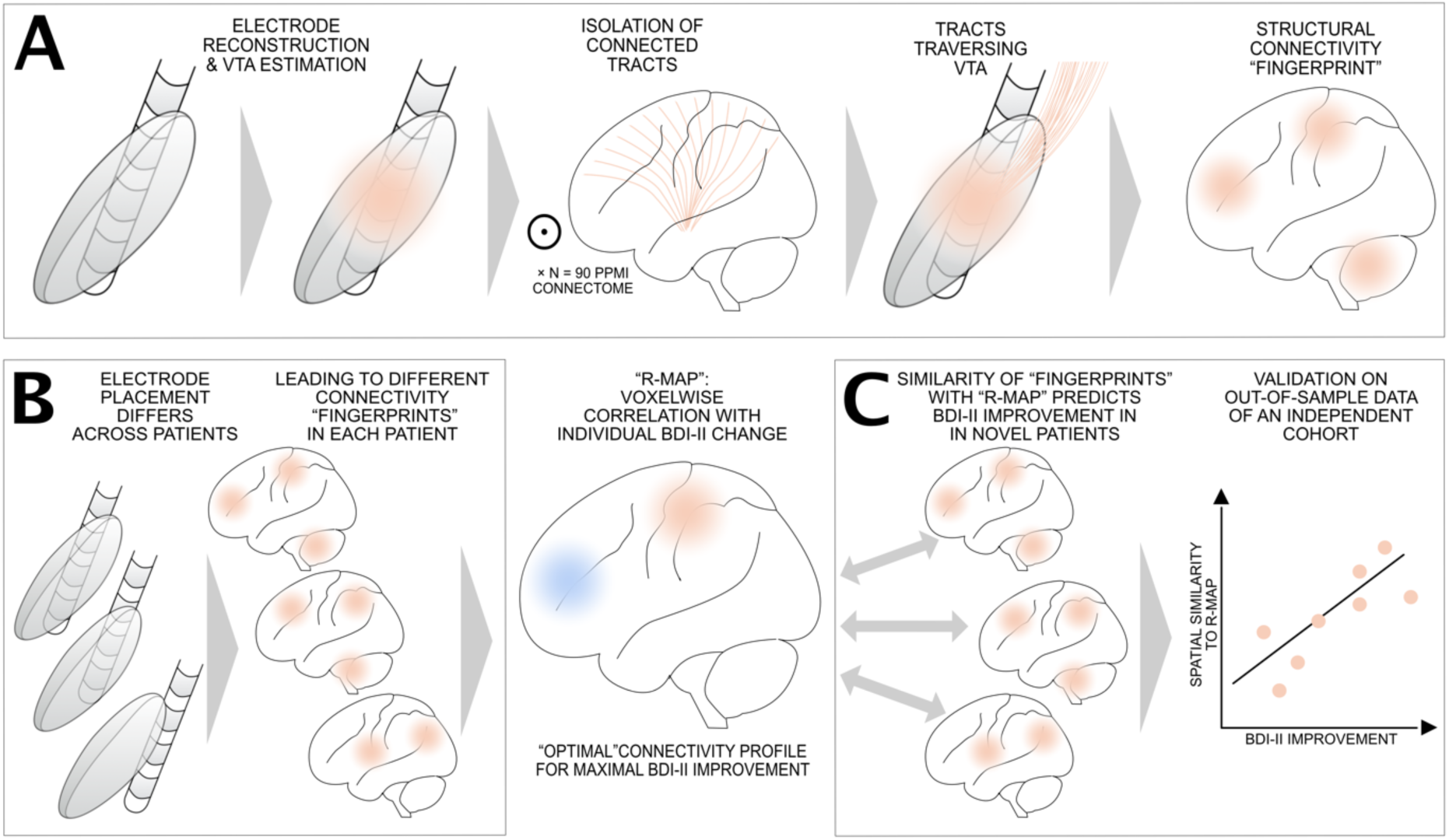
Overview of applied methods. A) In each patient, electrodes were localized and VTAs were calculated in standard stereotactic space using Lead-DBS software. From a normative Parkinson’s Disease connectome (N = 90 PPMI datasets), tracts that traversed through each patient’s VTA were selected and projected to the brain as fiber density maps. These maps represent the structural connectivity “fingerprint” seeding from each VTA. B) Varying electrode placement leads to different connectivity “fingerprints” in each patient. Across the group of patients, these fingerprints are used to generate a model of connectivity that is associated with maximal BDI-II improvement by voxel-wise correlation (“R-Map”). C) The R-Map represents a model that denotes how electrodes should be connected to result in maximal BDI-II improvement. When comparing each novel patient’s “fingerprint” with this model (by means of spatial correlation), individual BDI-II improvement can be predicted. Crucially, this is done to predict improvement in out-of-sample data, i.e. across cohorts or in a leave-one-out fashion throughout the manuscript. This means that the R-map is never informed by the predicted patient’s structural connectivity “fingerprint”.

Recently, it has been shown that using binarized VTAs (that would model all-or-nothing activations) could predict slightly less variance in clinical outcomes in comparison to using weighted VTAs such as the E-field gradient vector magnitudes^20^. Binary VTAs are based on specific thresholds that assume a certain type of axon diameter and orientation and do not grasp the anatomical complexity of the subcortex^25^. To account for this general limitation of the VTA concept, we repeated all analyses using the unthresholded E-field magnitude instead of the VTAs^26^.

Whole-brain structural connectivity profiles seeding from bilateral VTAs or E-Fields were estimated using a PD group connectome that is based on publicly available data (Parkinson’s Progression Markers Initiative; www.ppmi-info.org). This PPMI normative connectome of PD patients (n=90; age 61.38±10.42, 28 female) was priorly computed^24^ and has been used in context of DBS multiple times before^2,26,27^. For each patient, fibers passing through the VTA or a non-zero voxel of the E-Field were selected from this normative connectome and projected onto a voxelized volume in standard space (1mm isotropic resolution) while keeping count of the fibers traversing each voxel. In the VTA-based analyses, the number of fibers traversing each voxel was denoted (resulting in classical fiber-density map). In the E-Field-based analyses, each fiber received the weight of the maximal E-Field magnitude of its passage and fiber densities were weighted by these values.

### Modelling connectivity-driven mood changes

Structural connectivity strength, i.e. the number of fibers between VTA and each voxel was Spearman rank-correlated with BDI-II change (preoperative - postoperative), which resulted in a connectivity map that showed positive or negative associations with BDI-II improvement. In the following, these types of maps are referred to as R-maps (since they denote Spearman’s correlation coefficients for each voxel). Spearman’s correlation was used since tractography results are highly non-Gaussian distributed and rather follow an exponential distribution^28^. All analyses were carried out in Matlab (The Mathworks, Natwick, MA). We used randomized permutation tests (5000 permutations) to test for significance (at a 5% level) and used Spearman’s correlation coefficients throughout all analyses.

#### Validation of the training dataset

One R-map for each subset (BER, QU) was calculated. R-maps were then used to predict BDI-II changes in out-of-sample data (i.e. cross-predicting between QU ↔ BER cohorts) by spatial correlation between the R-map (model) and the connectivity profile seeding from the VTAs in each patient. This was done across voxels with an absolute Spearman’s R-value of >0.1 on each R-map. For example, the R-map (model) was calculated across the BER sample and voxels with an absolute R>0.1 were spatially correlated with connectivity maps in the QU sample. For each patient in the QU cohort, this led to one R-value that coded for spatial similarity to the model. These R-values were then correlated with empirical BDI-II changes. These head-to-head comparisons show direct relationships between BDI-II changes and the structural connectivity profiles of DBS electrodes. To further investigate their relationship with further clinical variables, the structural connectivity predictor was fed into regression models that included additional clinical information. This was done to analyse whether structural connectivity significantly explained variance above and beyond other clinical variables.

An additional leave-one-out cross-validation (i.e. data from patients 1-79 was used to predict patient 80 and so on) across the training sample (BER/QU combined) was run to test whether similarity to the specific structural connectivity profile of the training set (which is denoted by the R-map) could significantly predict absolute BDI-II change. Furthermore, we validated the results by running the analyses again (i) based on the E-field instead of VTA and (ii) using the percentage BDI-II change relative to baseline instead of the absolute BDI change. Moreover, to test for potential lateralization of connectivity profile, we reran analyses for left and right VTAs separately.

#### Prediction of the test dataset

In the same fashion as the cross-prediction between the sub-cohorts of the training dataset, a joint R-map for the entire training/cross-validation set (BER+QU) was generated, which was used to predict BDI-II change in patients of the test dataset (CGN).

#### Testing robustness of the model across the entire sample

We applied the leave-one-out cross-validation across the whole dataset, i.e. data from patients 1-115 was used to predict patient 116 and so on. Finally, to control for the effect of sex, postoperative LEDD, LEDD-DA and UPDRS-III reduction, those variables were included in the prediction models as covariates. In order to visualize possible confounding variables, we calculated the R-maps for subsamples of our patient cohorts: Figure 6 shows that the structural connectivity patterns predictive for postoperative BDI-II worsening remains in the left prefrontal cortex for A) patients that were not depressed before surgery (BDI-II<13, n = 86); B) patients that were mildly or moderately depressed before surgery (BDI-II >13, n = 30); and C) patients that were not treated with dopamine agonists before or after DBS surgery (n = 26).

### Isolation of fibertracts that are discriminative for mood changes

In an additional analysis, we sought to identify tracts that could discriminate patients with positive from negative BDI-II change. For each fibertract in the normative connectome (PPMI 90, see above), its accumulative E-Field vector magnitude while passing by each patient’s electrode was denoted. This value was then Spearman rank-correlated with each patient’s clinical change in depressive symptoms. Thus, a fibertract that passed close to active contacts of patients that had BDI-II improvement but far from active contacts in patients that had BDI-II worsening would receive a high Spearman’s R-value (and tracts exhibiting the inverse property received a highly negative R-value). These R-values were used to color-code fibertracts that were positively and negatively predictive of BDI-II improvement. This analysis was expected to show identical (or highly similar results) as the “R-map” method explained above but with the advantage of working on a tract-by-tract basis (instead of a voxel-wise fashion). Thus, it is ideal to visualize the actual fibertracts that were predictive of change in depressive symptoms.

## Results

### Clinical data

Disease duration in the entire sample (n=116; Table 1; Supplement 1) was 9.55±4.45 years. DBS lead placement was similar across all three cohorts (Fig. 2A,3C). Motor improvement with DBS was significant although measured ON medication reaching an average DBS response of 27.56±8.37% as measured by the UPDRS-III. Preoperative LEDD was 1142.46±567.47mg as compared to postoperative 464.45±291.37mg (56.31±29.71% reduction) with a contribution of dopamine agonists of 191.06±16.62mg pre- and 107.51±10.73mg postoperatively. Total LEDD, LEDD-DA, and UPDRS-III reduction were not significantly different in training and test datasets (*p*>0.05 for all three variables, Table 1). According to BDI-II scores prior to DBS surgery, 35 patients showed signs of mild depression, six of moderate depression but none were classified severely depressed (all BDI-II scores <29). Postoperative BDI-II assessments classified 31 patients as mildly depressed (of these twelve patients were classified not depressed prior to surgery), seven patients as moderately depressed (of these two patients had been mildly depressed before and two patients had been classified not depressed before surgery) and 2 patients as severely depressed (one had been classified as not depressed prior to surgery, the other had been moderately depressed). On average, BDI-II scores decreased from 9.94±5.41 to 8.96±6.43 (on average by 0.97±5.86 points) postoperatively, i.e. there was an overall reduction in BDI-II of 3.34±87.48% but the difference was not significant. Importantly, scores improved in 65 patients and worsened in 41 patients. Ten patients showed no change in BDI. Stimulation amplitudes between left and right hemispherical DBS electrodes (left: 2.46 ±0.79 mA; right: 2.58 ±0.83 mA) were not significantly different (p = 0.22).

**Figure 2:**
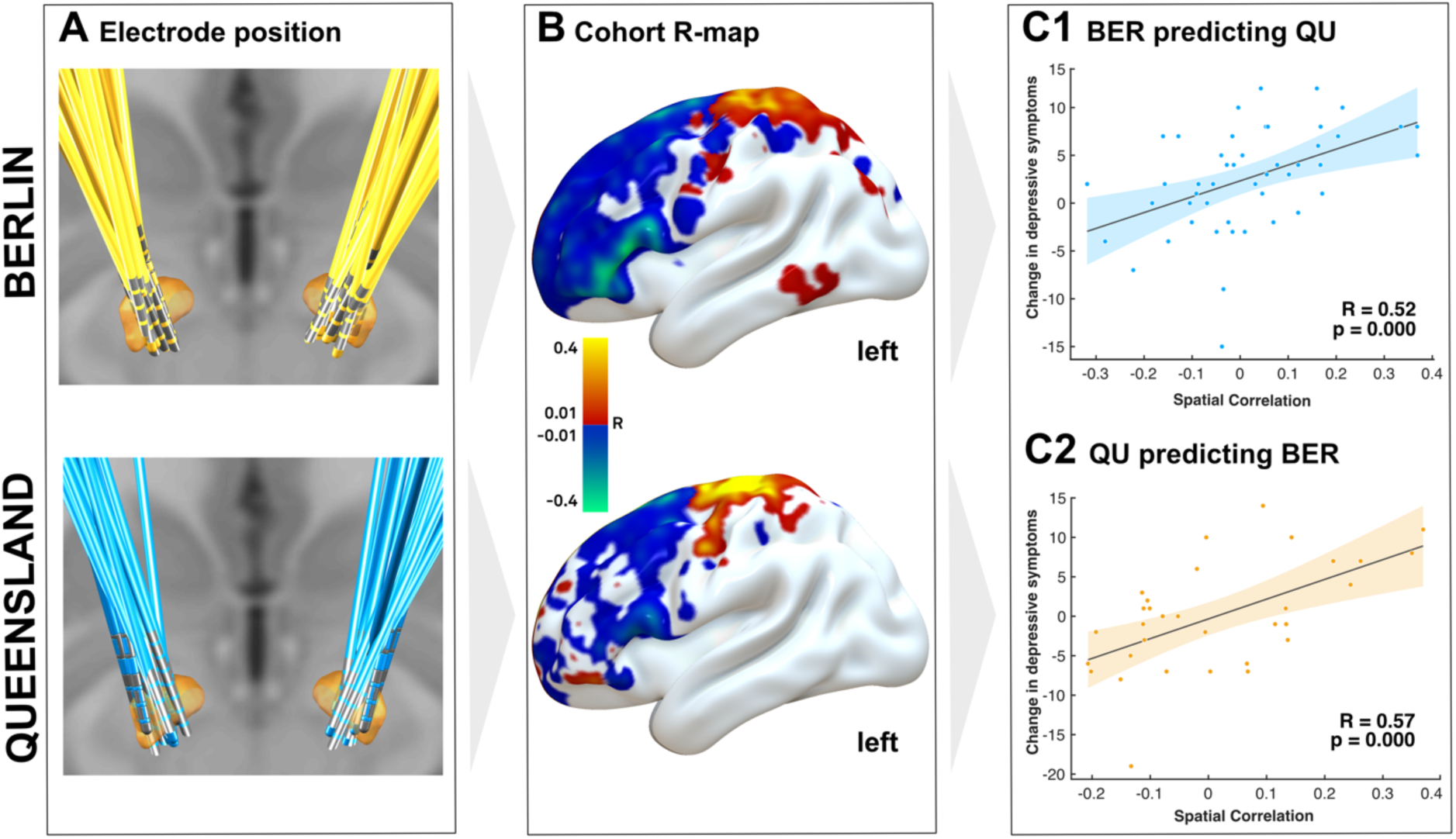
Structural connectivity predicting change in depressive symptoms in the training dataset (N = 80). A) Electrode position for the two cohorts from Berlin and Queensland. B) Each cohort’s R-Map represents the association with change in depressive symptoms under STN-DBS. Negative (blue) areas of the left hemisphere shown here relate to worsening of depressive symptoms. R-Maps revealed a significant association between worsening of depressive symptoms after STN-DBS and connectivity to left dorsolateral PFC. C1) Based on the R-Map from the Berlin cohort, depressive symptoms in the Queensland Cohort could be significantly predicted and vice versa (C2). R-Maps are presented smoothed with a 3mm full-width half-maximum Gaussian kernel to increase signal-to-noise ratio.

**Figure 3:**
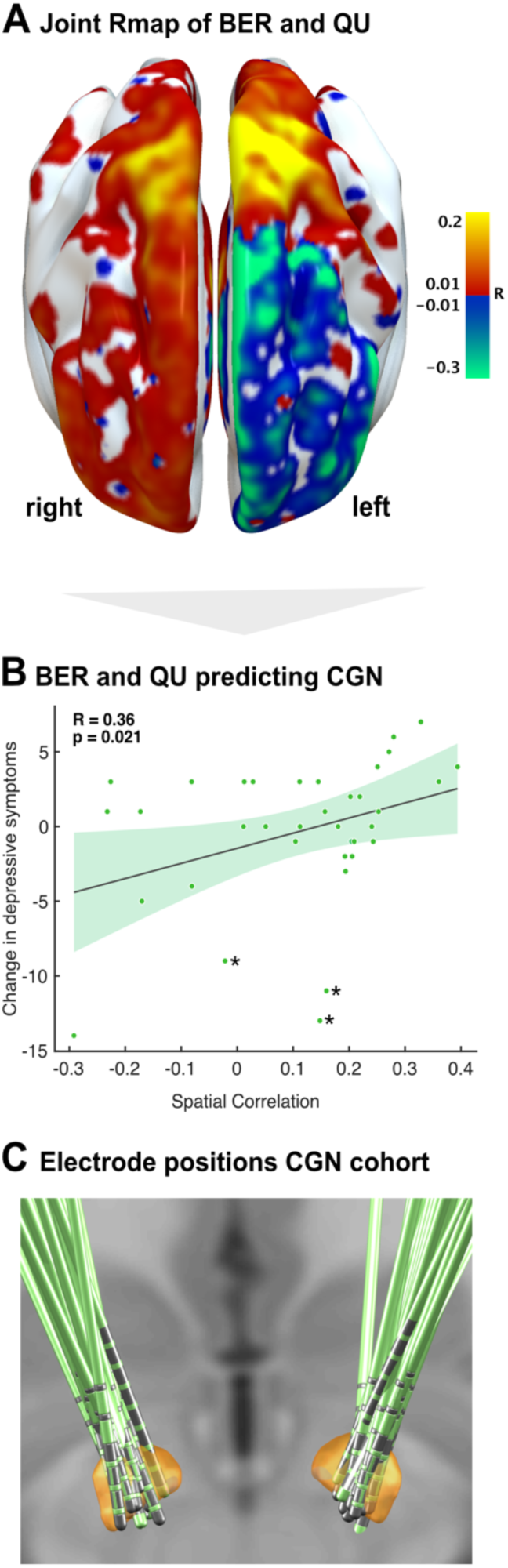
R-map of the training-dataset and prediction of the test-dataset. A) R-Map of the training dataset. Negative (blue) areas represent association with worsening of depressive symptoms while positive (red) areas represent association with improvement of depressive symptoms under STN-DBS. The R-Map is presented smoothed with a 3mm full-width half-maximum Gaussian kernel to increase signal-to-noise ratio. B) The R-Map of the training dataset (Berlin-Queensland model) significantly predicted change in depressive symptoms in the test-dataset (Cologne). C) Electrode positions of the test dataset within the STN.

### Connectivity related to DBS-induced mood changes

We identified a VTA-based structural connectivity map (R-map) predictive of postoperative BDI-II change in the training dataset (Fig. 3A). The more fibers connected a patient’s VTA to the positive areas (warm colors) of this map, the more their depressive symptoms improved postoperatively. On the contrary, the more a patient’s VTA was structurally connected to the negative areas (cold colors) of this map, the more their depressive symptoms worsened.

#### Validation of the training dataset

The R-maps of the two sub-cohorts in the training dataset were similar: On the right hemisphere of the R-map, connectivity to motor and prefrontal regions is universally associated with depressive symptom improvement. On the left hemisphere however, connectivity to the prefrontal cortex (PFC), including the dorsolateral PFC (dlPFC) is strongly associated with worsening of depressive symptoms, whereas connectivity to sensorimotor and superior parietal areas is associated with symptom improvement (Fig. 2). Cross-predictions were significant, i.e. the R-map based on BER-data could predict BDI-II change in relation to structural connectivity in the QU dataset (Fig. 2C, R=0.52, *p*<0.0001) and vice versa (R=0.57, *p*<0.0001). In a leave-one-out cross-validation across the training sample (BER/QU combined), similarity to this specific structural connectivity profile (R-map) could significantly predict absolute BDI-II change (R=0.26, *p*=0.01) even when basing structural connectivity profiles on the E-field instead of VTA (R=0.24, *p*=0.015) or when using the percentage BDI-II change relative to baseline instead of absolute BDI change (R=0.20, *p*=0.04). To test whether the effect was lateralized to either hemisphere, we reran analyses for left and right VTAs separately and found that connectivity on either hemisphere alone was predictive for BDI-II change (right: R=0.347, *p*=0.002; left: R=0.359, *p*=0.001). To rule out a systematic left-right bias of VTA placement, displacement of image centring or asymmetries in the normative connectome, we re-created the R-map after flipping the VTAs between left and right hemisphere while not flipping the connectome. The results remained stable, producing the same (lateralized, but mirrored) findings. This control analysis could largely rule out any systematic left-right bias in our analysis pipeline.

#### Prediction of the test dataset

The R-map based on the whole training set (BER/QU combined) was used to predict BDI-II change in the independent test-dataset (CGN) by calculating spatial similarity between each CGN-patient’s connectivity profile with the BER/QU R-map. This validated our results as a significant correlation was observed (Fig. 3B, R=0.36, *p*=0.012). The test-dataset (CGN) had been isolated from all model-selection processes and was used as a final confirmation after cross-validation between BER and QU. Still, the validation also yielded similarly positive predictions when using any other cohort as test-dataset (QU/CGN→BER: R=0.38, *p*=0.009; BER/CGN→QU: R=0.20, *p*=0.06). In the CGN test dataset some specific features were noted: patient #10 and #19 were diagnosed with comorbid depression and anxiety disorder at baseline; patient #13 reported pain and relatedly negative mood. Those three patients are marked with asterisk in Figure 3B. Importantly, excluding those patients still rendered the prediction significant (R = 0.28, *p* = 0.047).

#### Testing robustness of the model across the entire sample

As final step, we created one final R-map across all available data (BER/QU/CGN, n=116) which was predictive for BDI-II change in all three cohorts (Fig. 4A). This final R-map yielded significant predictions in a leave-one-out cross validation analysis (Fig. 4B, R=0.33, *p*<0.001). It remained significant when including sex, postoperative LEDD and LEDD-DA reduction and percentage UPDRS-III change (postoperative–preoperative ON medication) as covariates and correcting for cohort in a joint general linear model (R^2^=0.21, F_(112,104)_=3.88, *p*=0.001). Thus, this final model explained 21% of variance in BDI-II change based on clinical covariates and structural connectivity profiles across the whole group of subjects. The model remained significant when leaving any one cohort out (leaving out BER: R^2^=0.18, F_(82,76)_=3.25, *p*=0.01; leaving out QU: R^2^=0.29, F_(64,58)_=4.75, *p*=0.001; leaving out CGN: R^2^=0.20, F_(78,72)_=3.71, *p*=0.005). To test for the effect of UPDRS-III change induced by DBS (% difference OFF medication), a sub-analysis was performed for the entire BER sample (n = 32), where this data was available, and structural connectivity remained a significant predictor of BDI-II change (R^2^=0.19, *p*=0.05).

**Figure 4:**
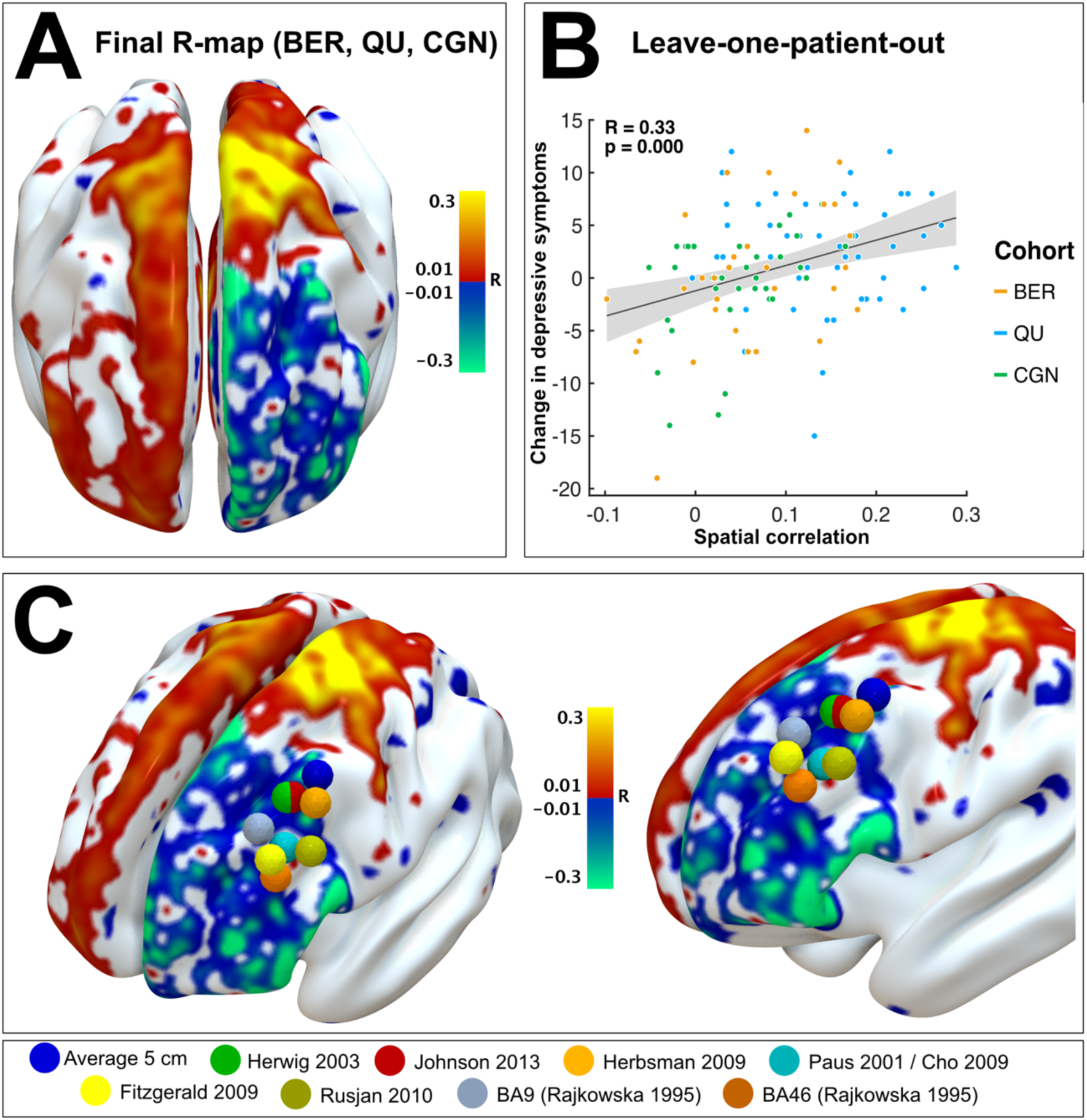
Final R-Map validation across all patients and proximity to TMS targets. A) R-Map associated with change of depressive symptoms over all patients (n = 116). B) Validation of the model using the approach of leaving-one-out design. C) rTMS targets for treatment of depression superimposed on final R-Map. R-Maps are presented smoothed with a 3mm full-width half-maximum Gaussian kernel to increase signal-to-noise ratio.

#### Focal impact on STN segments

To supplement the connectivity analyses, the number of voxels inside versus outside the STN and within motor versus nonmotor STN, that were stimulated by the VTAs were counted for each patient and correlated with BDI change. The more VTA-voxels lay inside the STN, the higher BDI-II scores improved (R=0.285, p=0.002), no matter if VTA stimulated motor or nonmotor STN segments (R=0.267, *p*=0.002 and R=0.197, *p*=0.034). The number of VTA-voxels outside the STN was not predictive for BDI-II change (R=-0.129, *p*=0.166), suggesting that stimulation of specific fibers passing the STN would explain BDI-II worsening, rather than unspecific stimulation of non-STN tissue.

#### Individual differences

We demonstrate that the connectivity profile shown in figure 4 was associated with changes on the BDI-II score across DBS centers, surgeons and cohorts. However, on a single patient level, changes showed a root-mean-square (RMS) error of 4.11±3.51 points in the leave-one-out cross-validation.

### Fibertracts related to mood changes

An additional analysis was run to identify the actual tracts (instead of their cortical projection sites) the modulation of which was correlated with BDI-II improvement. This was done on a tract-by-tract instead of voxel-wise basis but further confirmed our results using a different analysis pathway. Crucially, this data-driven analysis revealed largely more tracts on the left hemisphere than on the right hemisphere, again suggesting an impact of left DBS stimulation on change of depressive symptoms (Fig. 5A). Using lower thresholds, the pattern was similar between the two hemispheres but left hemispheric tracts were more predictive of BDI-II change and predictive tracts were found in larger quantities. The analysis revealed that the positively and negatively associated tracts seemed to differ in their anatomical course in that the negatively associated tract passed by the STN medial and at level of its limbic/associative functional zone, while the positively correlated tract passed through and slightly lateral to the motor STN (Fig. 5B). Moreover, as can be seen in Fig. 5C, the negatively associated tract traverses more laterally when ascending to the PFC.

**Figure 5:**
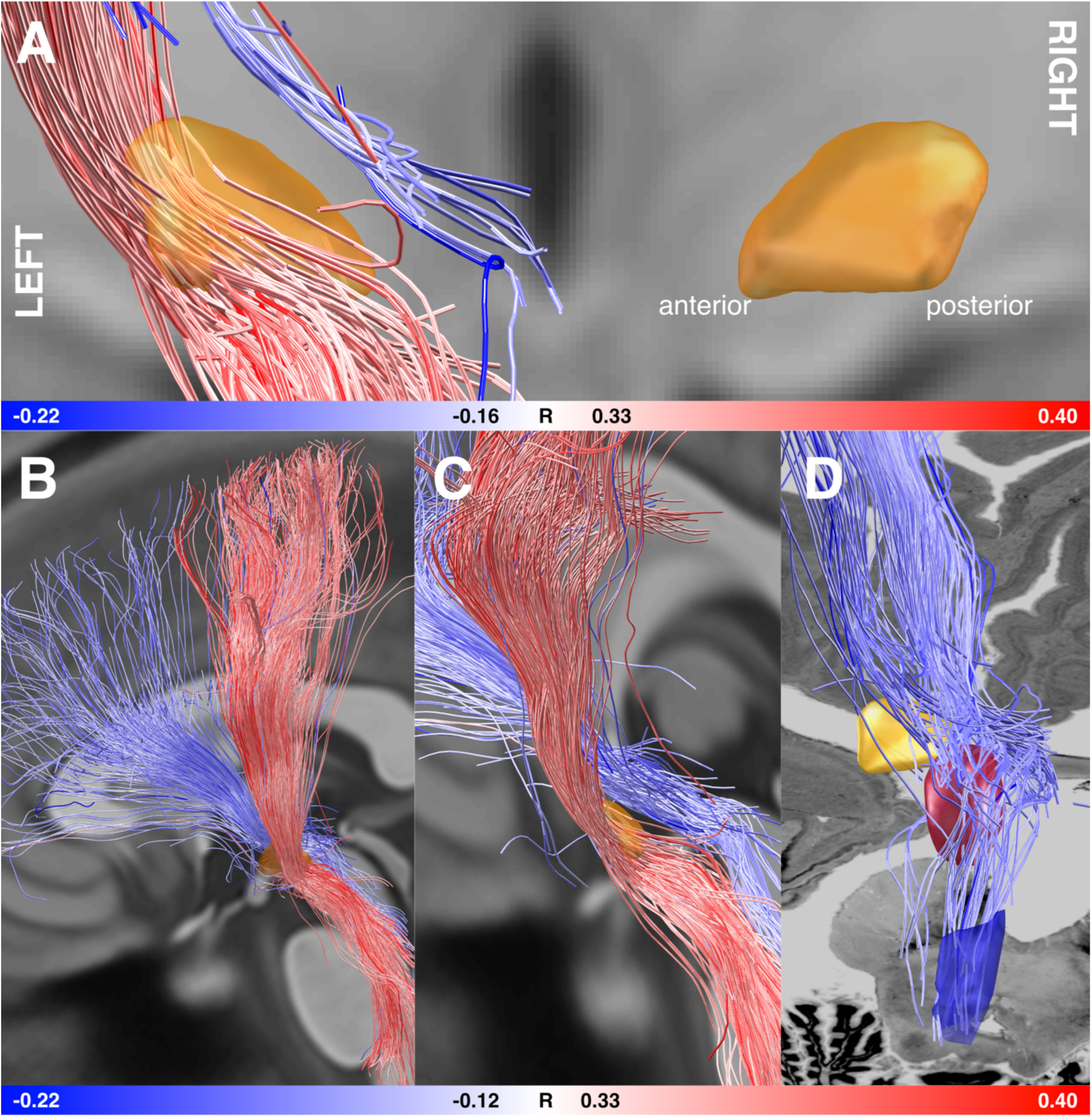
Fibertracts discriminative of BDI-II improvement when modulated. Red tracts are positively, blue tracts negatively correlated with clinical improvement. STN shown in orange. A) Coronal view from posterior with both hemispheres. At this threshold level, no fibers on the right hemisphere were associated with clinical improvement but a strong set of both positive and negative fibers were found on the left hemisphere. B) View from the left and C) view parallel to the longitudinal axis of the left STN. Positively and negatively correlated fibertracts seem to be distinct tracts, the positive one passing through the STN and lateral to it, the negative one medial and anteriorly. D) Superimposed on a section of the BigBrain ultrahigh resolution human brain model^56^, at the level of the brainstem, the negative tract seems to traverse around the red nucleus and may connect to (or originate from) brainstem nuclei such as the left dorsal raphe nucleus (shown in dark blue as defined by the Harvard Ascending Arousal Network Atlas).

**Figure 6:**
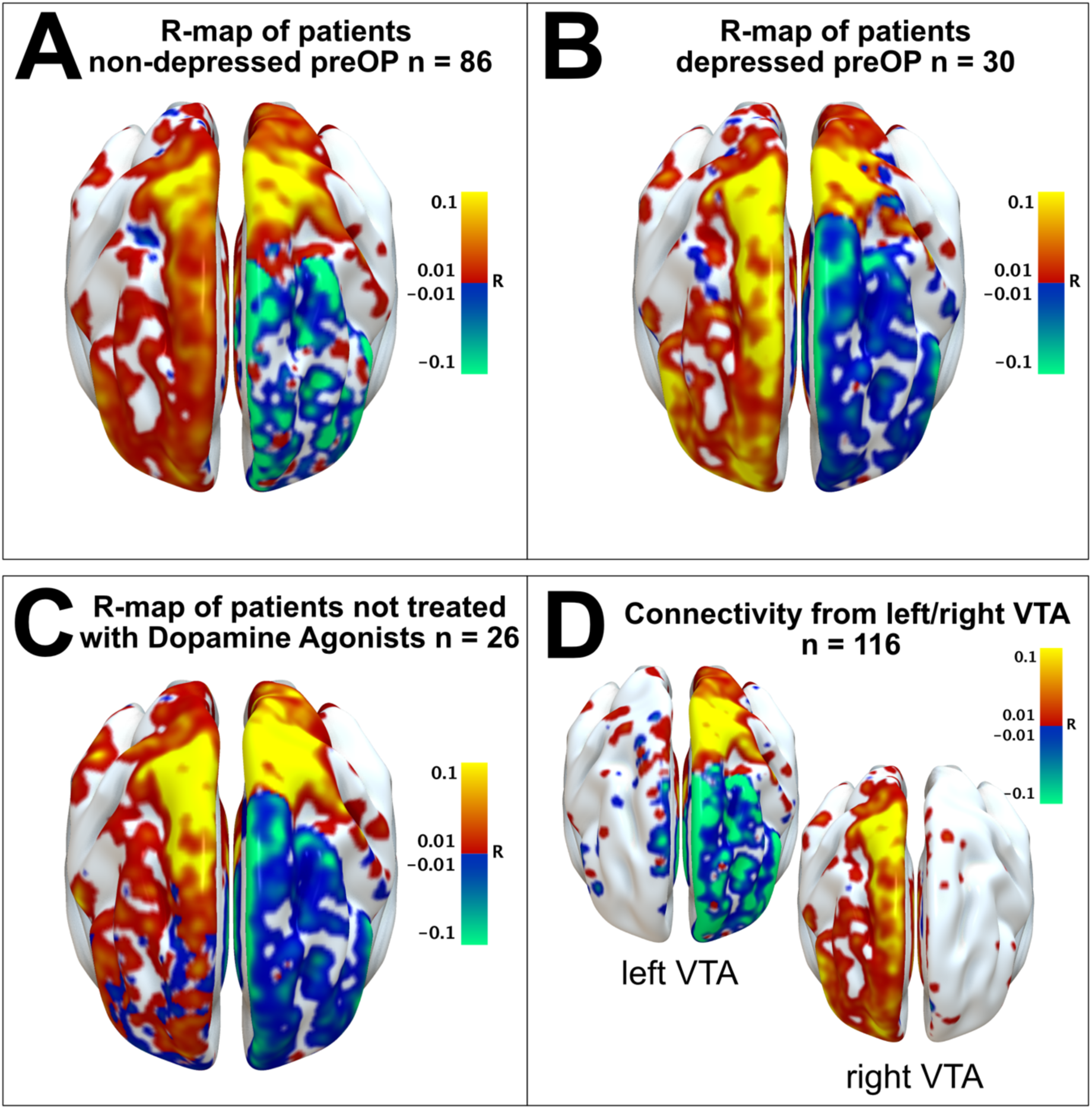
R-maps of subsamples of PD patients. The structural connectivity pattern that is predictive for BDI-II change remains stable in sub-analyses on A) patients that were not depressed before surgery (BDI-II<13, n = 86); B) patients that were mildly or moderately depressed before surgery (BDI-II >13, n = 30); and C) patients that were not treated with dopamine agonists before or after DBS surgery (n = 26). D) When carrying out the analyses for left and right VTAs separately, the connectivity pattern also exhibits the same spatial profile.

## Discussion

In this study, we modelled structural connectivity predicting changes in depressive symptoms following STN-DBS in PD. We identified a distinct connectivity pattern linking worsening of depressive symptoms under STN-DBS to left prefrontal impact. This connectivity profile predicted data across cohorts. Specifically, left-hemispheric impact on fibers anteromedial to the STN connecting to the PFC predicts worsening of depressive symptoms.

A common assumption on DBS-induced affective changes is their relation to rapid withdrawal of dopaminergic replacement therapy after surgery which increases anhedonia induced through affective network dysregulation^14^. While this factor explains acute and subacute postoperative affective changes, in our large multi-center sample, LEDD and LEDD-DA reduction did not explain BDI-II change after 6-12 months. Perhaps this relates to clinicians addressing this potential risk-factor for depression during long-term follow-up. Others have also reported a lack of correlation between LEDD reduction and non-motor PD symptoms like apathy and mood^6,8^. Yet, the general notion is that STN-DBS mimics the action of dopaminergic agents^10^ and acute stimulation more likely leads to hypomania than depression^10,29,30^. Interestingly, long-term improvement in motor symptoms supposedly inducing secondary improvements of mood^2^ did not change the predictive value of connectivity for depressive symptom change, suggesting stimulation influences affective processing more directly, i.e. via impact on left PFC.

The association of depressive symptoms and connectivity to the left PFC is unsurprising given the vast amount of evidence linking depression to frontal lesions: Hypoactivity and dysfunction of left PFC is commonly found in patients with depression^31^. Moreover, depressive symptoms increase gradually after left dlPFC traumatic brain injury and stroke depending on the extent of damage and network impact^32^. Indeed, large-scale network effects, hemispheric asymmetries and connectivity play an important role in the development of depressive symptoms^33^; e.g. post-stroke depression has been linked to altered dlPFC functional connectivity^34^. Functionally, the left dlPFC might regulate negative affect via the frontoparietal cognitive control network^35^. In major depression, excitability of hypoactive dlPFC tissue^31^ is augmented with non-invasive high-frequency repetitive transcranial magnet stimulation (rTMS) leading to symptom amelioration^36^. Although the precise mechanism of dlPFC rTMS in improving depressive symptoms is not fully understood, a role of local and remote network changes and altered PFC connectivity is evident^37^. Interestingly, common targets of rTMS in depression^37^ precisely lie within the clusters we find associated with BDI-II worsening under STN-DBS; Fig.4C). Thus, worsening of depressive symptoms following STN-DBS might occur due to tempering of fibers linked to the left dlPFC.

Concurrently, in this study worsening of depressive symptoms under STN-DBS is associated with stimulation of fibers connecting prefrontal areas via zona incerta to the dorsal mesencephalon and brainstem (Fig.5). We presume that STN-DBS disrupts information flow along these fibers. One candidate brainstem region whose link to the PFC might be disturbed by DBS leading to depression is the dorsal raphe nucleus (DRN), which as part of the serotonergic system is known to impact mood states and which is hypoactive in depression^38^. Indeed, unbalanced prefrontal-DRN connectivity relates to depression^39^ and rodent studies have shown that STN-DBS may inhibit serotonergic output from the DRN inducing depression-like behavior^40^. Since there are no direct STN-DRN connections, misplaced STN-DBS may accidentally disrupt left prefrontal-DRN serotonergic communication^41^, indirectly fostering depressive states. Another candidate neural substrate involved is the ventral tegmental area, which as origin of mesolimbic dopamine projections is pivotal for reward-processing but also plays a role in depression^42^. Yet, this neural substrate is less likely given the exact anatomical course of the tract.

As mentioned in the introduction, STN-DBS has led to improvements, worsening or no effect on affective symptoms when analysed on a group level. The same applies to the combined cohort analysed here. While depressive symptoms in individuals of the cohort improved and got worse – and for some by large – across the total cohort, preoperative and postoperative scores were not significantly different. On a center-by-center level, one cohort (QU) significantly improved while the other two did not. Our study may provide a reason for these conflicting results which could be based on electrode placements, especially in the left STN. A left electrode position that is more anterior was associated with worsening of affective symptoms. Crucially, this was the case in all three cohorts. Thus, depending on surgical practices or slightly differing average targets of each center, differing group effects in reported studies or DBS cohorts could be explained.

Although activation of fibertracts bypassing the STN explained *worsening* of depressive symptoms, a role of the STN itself in affective processing is undisputed^17^. Like the striatum, the STN is a node of convergence of affective, cognitive and motor input^4,43^. Its activity is modulated through coupling with PFC activity^44^ and STN-DBS impacts affective processing^45^. Concurrently, in our data depressive symptoms *improved* if predominantly fibers connecting the dorsolateral (motor) STN with the sensorimotor cortex were stimulated (as reported previously^46^). Although STN-sensorimotor-cortex connectivity has primarily been associated with improvements in *motor* performances a recent study showed mood improvement actually relies on DBS of the motor STN^47^. Similarly, motor STN stimulation can normalize cognitive performance^48^. This implicates the overlapping presence of affective/associative and motor processing neurons in the STN motor segment^4^ and suggests that retuning the motor loop may improve non-motor features as well.

Another region to which electrode connectivity predicted improvement of depressive symptoms was the right dlPFC. Current neuroimaging studies speak of reduced left and increased right dlPFC activity associated with depressive symptoms^49^. Although this fits our results, we must emphasize that while evidence on the link of a *left hypoactive* dlPFC with depressive symptoms is solid, the association of *right hyperactive* dlPFC with improved mood seems still weak^50^ and the origin and validity of such laterality effects require further investigation.

Taken together, in the left hemisphere, high-frequency stimulation of fibers anteromedial to the STN is associated with worsening of depressive symptoms while stimulation of dorsolateral STN leads to improvement of depressive symptoms in PD patients. The connectivity profile described in this study may inform surgeons and clinicians in placement and settings of STN-DBS. The present results may help avoid harmful side effects of STN-DBS in PD patients by considering connectivity to networks guiding these side effects, however further work fostering a refined understanding of the functionality of PFC-STN connectivity and lateralized impact is key.

As a final consideration, it is important to stress that we believe depression is a system-level disorder: no single brain region or neurotransmitter is the sole driving force but instead, integrated cortical-subcortical networks seem to be key^31^. This means the impact of STN-DBS on affective networks based on patients’ connectivity profiles is surely not the only factor contributing to changes in depressive symptoms. Yet, this research may contribute to better understand, avoid and treat affective side effects like depressive symptoms in patients with STN-DBS.

There are several limitations to our findings. First, differences in the timepoints of assessment of depressive symptoms across DBS centers might have affected our results. There is variance in follow-up timepoint between cohorts which arose given the retrospective multi-center study-design. However, all follow-ups were later than 5 months after surgery when usually a stable effect of DBS parameter settings and medication is reached. Moreover, this variance might have biased our results toward non-significance and could also be interpreted as a strength in robustness of results. We do believe though that with our large sample size slight variances in the timepoint of BDI-II assessment did not systematically bias our results.

Second, there is a variation in electrode type in the patients included in this study. This could affect the VTA-model, e.g. by respective consideration of constant voltage versus constant current default settings in DBS systems. To circumvent a bias of this factor, we reran analyses using the unthresholded E-field surrounding electrodes and found similar results.

Third, we used an age- and disease-matched group connectome that was based on data acquired in PD-patients enrolled in the PPMI project, which will not account for individual structural connectivity but instead assumes similar connectivity profiles in all patients. While this assumption might not hold true in all cases, several recent studies have introduced and validated the method within DBS context^2,3,27^. Beyond practical advantages (where patient-specific connectivity data is often not available and cannot be acquired postoperatively), normative connectomes like the one used here comprise a high N of subjects which by averaging lead to high signal-to-noise levels and state-of-the-art data quality. Yet, residual movement during the scan might be a potential source of confound on the data quality of the images which remains a limitation. Moreover, disease-stage of patients in the PPMI-connectome differed from our patients in that they were predominantly early-stage PD patients. While prior research showed that the choice of group connectomes does not largely alter results in the kind of analyses performed here^2^ we still emphasize this limitation.

Fourth, we only had UPDRS-III scores *ON medication* for most patients and thus pre- to postoperative comparisons might not reflect the full impact of STN-DBS on motor symptoms.

However, a sub-analysis of the entire BER sample (n= 32) including UPDRS-III comparison *OFF medication* yielded similar results; thus, we believe that structural connectivity remained the strongest predictor for depressive symptom change.

Fifth, we used a general self-rating scale to score depressive symptoms, which does not allow to specify the change in subitems such as anxiety or apathy. Unfortunately, our study was unable to address DBS-related changes in symptom-specific pattern since the corresponding scores were not available in our cohorts. Thus, future research should compare our results with other symptom clusters.

Sixth, a confound of the present study should be seen in the accuracy of electrode reconstructions. While it is impossible to estimate an empirical average error without histological postmortem validation of results, we applied a modern pipeline that was explicitly designed for the task. Processing steps included phantom-validated electrode reconstructions^51^, brain shift correction^20^, multispectral nonlinear registrations to target nuclei that were empirically validated^22^ and finite-element method-based estimations of stimulation volumes^2^. Two studies have validated the pipeline using electrophysiological recordings^52,53^ and others have demonstrated that results are capable of explaining clinical improvements across patients, DBS centers and cohorts^2,3,48,54,55^. Still, residual errors of electrode reconstructions cannot be excluded.

Finally, while the connectivity profile associated with depressive symptom changes identified in this study could be reproduced in three individual cohorts and cross-validation across DBS centers yielded significant predictions, on an individual level, predictions are not truly useful for clinical practice with an average RMS error of 4.11±3.51. Thus, our model is not yet able to predict clinical changes along symptoms covered by the BDI-II score with high precision. While this is a limitation of clinical applicability, the main message of our results may still be useful to inform clinical practice, especially in targeting the left electrode for STN-DBS.

In conclusion, the present results have potential therapeutic value for the refinement of brain stimulation targets. In personalized brain stimulation, identifying proximity to fibers connecting the electrode with the left dlPFC might have prognostic utility in predicting affective changes under STN-DBS. Prospectively, connectivity maps as well as isolated fibertracts can be used in surgical planning to optimize positioning of DBS leads. Furthermore, with directional leads, the electrical field could be guided away from fibertracts anteromedial to the left STN to depressive symptom worsening. Importantly, this study specifically shows that the STN connectivity profiles might have to be treated differently for the right and left hemisphere.

However, more work is needed to validate this presumption in patient-specific connectivity. Altogether, our findings contribute to understanding of how negative mood effects may originate following STN-DBS and pave the way toward personalized brain stimulation in which individual connectivity profiles and symptom constellations determine optimal DBS targets.

## Acknowledgements

This study was supported by the Deutsche Forschungsgemeinschaft (DFG grant SPP2041, “Clinical connectomics: a network approach to deep brainstimulation” to A.A.K. as well as Emmy Noether Grant 410169619 to A.H.).

We thank Alexandra Horn for her valuable contribution in the data analysis.

Data used in the preparation of this article were obtained from the Parkinson’s Progression Markers Initiative (PPMI) database (www.ppmi-info.org/data). For up-to-date information on the study, visit www.ppmi-info.org. PPMI - a public-private partnership - is funded by the Michael J. Fox Foundation for Parkinson’s Research and funding partners, see www.ppmi-info.org/fundingpartners.

## Author contributions

F.I., A.H. and A.A.K. contributed in the design of the study. All authors contributed to acquisition and analysis of data. F.I., A.H. and A.A.K. drafted the text and prepared the figures. All authors reviewed and revised the manuscript for intellectual content.

